# Multiple constrained minimum variance beamformer (MCMV) performance in connectivity analyses

**DOI:** 10.1101/567768

**Authors:** Adonay S. Nunes, Alexander Moiseev, Nataliia Kozhemiako, Teresa Cheung, Urs Ribary, Sam M. Doesburg

## Abstract

Functional brain connectivity is increasingly being seen as critical for cognition, perception and motor control.Magnetoencephalography and electroencephalography are modalities that offer noninvasive mapping of electrophysiological interactions among brain regions, yet suffer from signal leakage and signal cancellation when estimating brain activity. This leads to biased connectivity values which complicate interpretation. In this study, we test the hypothesis that a Multiple Constrained Minimum Variance beamformer (MCMV) outperforms the more traditional Linearly Constrained Minimum Variance beamformer (LCMV) for estimation of electrophysiological connectivity. To this end, MCMV and LCMV performance is compared in task related analyses with both simulated data and human MEG recordings of visual steady state signals, and in resting state analyses with simulated data and human MEG data of 89 subjects. In task related scenarios connectivity was estimated using coherence and phase locking values, whereas envelope correlations were used for the resting state data. We also introduce a novel Augmented Pairwise MCMV (APW-MCMV) approach for signal leakage suppression in resting state analyses and assess its performance against LCMV and more conventional MCMV approaches. We demonstrate that with MCMV effects of signal mixing and coherent source cancellation are greatly reduced in both task related and resting state conditions, while in contrast to other approaches 0-and short time lag interactions are preserved. In addition, we demonstrate that in resting state analyses, APW-MCMV strongly reduces spurious connections while better controlling for false negatives compared to more conservative measures such as symmetrical orthogonalization.

## Introduction

Since Biswal et al. discovered in 1995 that the sensorimotor cortex activity at rest was correlated with its interhemispheric counterpart, neuroimaging of functional brain connectivity has been the focus of increasing research interest. Brain connectivity has been critical for understanding the intrinsic organization of brain function at rest (Fox et al., 2005; de Pasquale et al., 2010; Yeo, Krienen, & Sepulcre, 2011; van den Heuvel & Sporns, 2013), as well as how different task demands reshape the organization of large-scale brain connectivity (Krienen, Yeo, & Buckner, 2014; Liljeström, Stevenson, Kujala, & Salmelin, 2015; O’Neill, et al., 2017; Saarinen, Jalava, Kujala, Stevenson, & Salmelin, 2015; Ribary et al. 2017). Mounting evidence indicates specific functional connectivity alterations in various pathologies of the nervous system (Lynall et al., 2010; Schoonheim et al., 2013; Bozzali et al., 2015; Cerliani et al., 2015; Pang & Snead III, 2016), and adequate methods for studying interactions among brain regions in function and dysfunction is of critical importance.

Electroencephalography (EEG) and magnetoencephalography (MEG) are modalities very well suited to study brain connectivity dynamics due to their high temporal resolution. To estimate source-space brain dynamics, however, it is necessary to reconstruct the activity at desired brain locations from sensor data located outside the head. One popular inverse model method is linearly constrained minimum variance (LCMV) beamforming (Van Veen, van Drongelen, Yuchtman, & Suzuki, 1997; Vrba & Robinson, 2001; Kensuke Sekihara, Nagarajan, Poeppel, Marantz, & Miyashita, 2002; S E Robinson, 2004; K. Sekihara et al., 2007; Herdman & Cheyne, 2009; Moiseev & Herdman, 2013) which has been shown to be very reliable in localizing sources and estimating brain activity (Kensuke Sekihara et al., 2002; Dalal et al., 2008; Murzin, Fuchs, & Kelso, 2011; Jonmohamadi et al., 2014). Inverse modeling, however, is an ill posed problem (Michel et al., 2004; Greenblatt, Ossadtchi, & Pflieger, 2005). Its solutions typically exhibit a degree of linear mixing of the estimated brain sources, especially for nearby sources. Connectivity measures sensitive to linear mixing or signal leakage are severely confounded as spurious interactions caused by linear mixing are difficult to distinguish from genuine interactions and this limits the usability and interpretation of the results (Schoffelen & Gross, 2009).

One way to overcome signal leakage is to use connectivity measures which discard interactions with zero time lags, such as phase lag index (Stam, Nolte, & Daffertshofer, 2007), imaginary coherence (Nolte et al., 2004) and orthogonalized envelope correlations (Brookes, Woolrich, & Barnes, 2012; Hipp, Hawellek, Corbetta, Siegel, & Engel, 2012). However this is a conservative approach that discards not only signal leakage within the zero phase lag interaction but genuine neurophysiological interactions with small time lags whose existence has been shown to reflect real neural activity by invasive brain recording studies (Brody, 1999; Lachaux, Rodriguez, Martinerie, & Varela, 1999; Lutz, Lachaux, Martinerie, & Varela, 2002; Singer, 2010). Furthermore, although such measures discard direct zero-lagged interactions, they are still susceptible to secondary leakage caused by a third source leaking onto the two tested sources at a non-zero phase lag. Although secondary leakage is much smaller, it is still an important confound that renders inaccurate estimates of connectivity (Palva & Palva, 2012; Wang et al., 2018).

Multiple Constrained Minimum Variance (MCMV) has been proposed as an alternative approach for signal leakage suppression without discarding zero-lagged interactions (Dalal, Sekihara, & Nagarajan, 2006; Popescu, Popescu, Chan, Blunt, & Lewine, 2008; Hui, Pantazis, Bressler, & Leahy, 2010; Moiseev, Gaspar, Schneider, & Herdman, 2011; Quraan & Cheyne, 2010). In this study, we will use an approach introduced in Moiseev et al. 2011 for localizing and estimating time courses. This type of MCMV is a multi-source variant of a scalar beamformer (Stephen E. Robinson & Vrba, 1999). We will use the term LCMV to refer to a traditional single source scalar beamformer. Beamformers work by constructing a spatial filter (or weights) for a given location that optimally suppresses interference from other locations. MCMV differs from LCMV in that it adds special constraints explicitly preventing linear mixing of reconstructed sources. Rather than orthogonalizing the reconstructed sources time courses, this approach orthogonalizes the weights of one location with the forward solutions of all the other sources included in the beamformer (albeit, the number of sources is limited to a few). Consequently, the activity in other locations does not influence the estimation of the activity of a target location, dealing, then, with the signal leakage problem and coherent source cancellation.

Previous studies (Moiseev et al., 2011; Herdman, Moisev, & Ribary, 2018) introduced unbiased MCMV estimators of source locations and orientations and demonstrated improved performance of MCMV compared to LCMV in reconstructing source time courses. The application of MCMV to connectivity analyses, however, has yet to be thoroughly characterized. The aim of this study is to investigate the performance of MCMV in connectivity analyses, relative to LCMV, under task and rest conditions. An important difference is that in task analyses, a noise covariance can be reliably estimated using inter-trial periods, whereas in resting state analyses those do not exist. This, in addition to poor signal-to-noise ratios, makes resting state connectivity estimates much more difficult and require different approaches, as described below. In the task-related analyses connectivity was estimated using coherence and phase locking values (PLV). First (i), a simple sinusoidal simulation was used to demonstrate spurious connectivity arising from signal leakage and signal cancellation effects. Second (ii), signals from ECoG, similar in complexity as M/EEG signals, were used as ground truth for the simulations. Third (iii), real human MEG recordings of steady-state foveal stimulation task were used to estimate connectivity in entrained areas. For all resting state analyses, envelope correlation was chosen as a connectivity measure due to its demonstrated reliability (Colclough et al., 2016) and popularity in resting state MEG studies. Source reconstruction was done using LCMV, a pair-wise MCMV (PW-MCMV) and an augmented pair-wise MCMV (APW-MCMV). We also used Symmetrical Orthogonalization (SO) applied to the LCMV time series (Colclough et al. 2015), as a reference approach for comparison. Note that this method involves discarding zero-lag interactions. In resting state analyses, first (i) data from a single subject was used to estimate connectivity and construct a simulation where the ground truth was known; second (ii), connectivity in real data was estimated at a group level; and third (iii), surrogate data without genuine connectivity was used to compare group level spurious connectivity (if any) reported by LCMV, MCMV and SO-based methods.

## Methods

In this manuscript, the following conventions will be used. Scalar quantities and components of matrices and vectors will be denoted by normal lowercase letters, vectors – by bold lowercase letters, and matrices – by bold capital letters. A superscript “T” denotes matrix transposition, and angular brackets 〈…〉 – statistical averaging.

### LCMV spatial filter

In the LCMV approach, reconstructed brain signal *s*_***q***_(*t*) at time *t* is represented as:

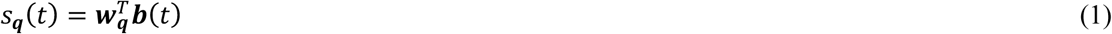

Here, parameter set ***q*** = {***r***, ***u***} consists of source location ***r*** and its orientation ***u***, the latter being a unit vector; ***b***(*t*) is a (*M × 1*) column vector of sensor readings at time *t* and *M* is a total number of sensors in EEG/MEG array; ***w***_***q***_ is a (*M × 1*) vector of LCMV filter weights. Weight ***w***_*q*_ is selected so that to minimize average reconstructed source power 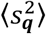 subject to a unit gain constraint 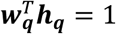, where ***h***_***q***_ denotes a (*M × 1*) forward solution vector for the source with parameters ***q*** = {***r***, ***u***}.

Assuming that ***b***(*t*) is a stationary random process with a zero mean, a well-known result is that optimal filter weights ***w***_***q***_ are given by the expression (Frost, 1972; Van Veen, et al., 1997; K. Sekihara, et al., 2007;):

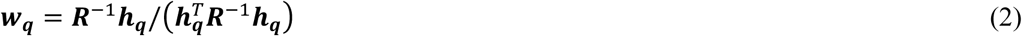

where ***R*** = 〈 ***bb***^T^ 〉 is the covariance matrix of the sensor data.

### MCMV spatial filter

The MCMV filter generalizes LCMV by reconstructing simultaneously *n* sources *s*_***q**i*_, *i* = 1, …, n. Specifically, a (n × 1) vector of source amplitudes ***s*** = {*s*_***q**1*_, …, *s*_***q**n*_} is represented as

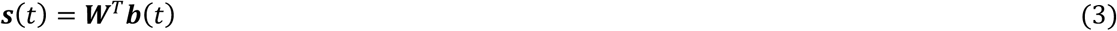

with ***W*** being a (M × n) matrix of weight vectors corresponding to each source: ***W*** = {***w***_***q**1*_, …, *w*_***q**n*_}. Weights ***W*** are selected so that to minimize *total* average reconstructed source power 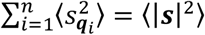 subject to a constraint ***W***^T^***H*** = ***I***_*n*_. Here we defined a (M × n) joint forward solution matrix ***H*** = {***h***_***q**1*_, …, *h*_***q**n*_}, and ***I***_*n*_ is an n-dimensional identity matrix. The weight matrix which satisfies the minimum variance, i.e. minimum power, is (Frost, 1972; Van Veen, et al., 1997; K. Sekihara, et al., 2007):

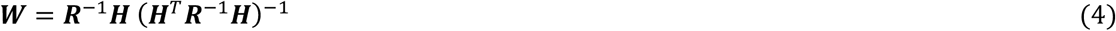

Importantly, due to constraints ***W***^T^***H*** = ***I***_*n*_, each weight vector ***w***_***q**i*_ satisfies a “unit gain condition” with respect to its own forward solution: 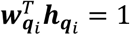, and a “zero gain” condition with respect to all other sources: 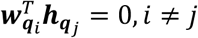. The latter property makes MCMV beamformer insensitive to source cancellation effects due to source correlations, and prevents linear mixing of reconstructed source time courses.

In MCMV reconstruction, the following approach was used to define the beamformer order. For task-related source estimation, where a limited set of sources of interest could be identified, the order was equal to *n* = number of sources. For resting state connectivity analysis, we estimated sources using a) a pair-wise MCMV (PW-MCMV) using 2-source MCMV beamformer so that ***s***_*pair*_ = {*s*_***q***1_, *s*_***q***2_}, thus preventing direct mixing of the sources in the pair, b) an augmented pair-wise MCMV (APW-MCMV) method, detailed below, where in addition to the two sources of interest, we included up to two more sources from locations adjacent to each source of interest, so the maximum beamformer order could reach up to six. The goal was to reduce the possibility of false connectivity in the pair due to *indirect* leakage.

### Source localization

In this section, we give a brief summary of how MCMV and LCMV source localization was performed. Regarding the MCMV approach, details and derivations may be found in Moiseev et al. 2011. LCMV has been used in many studies; a comprehensive analysis and review of relevant literature can be found in (K Sekihara & Nagarajan, 2008).

In the beamformer approach, to find the sources of brain activity special localizer functions (“localizers”) defined on a source parameter space are used. The latter includes 3D positions and orientations for each source and therefore has *5n* dimensions, where *n* is the beamformer order. The maxima of the localizer points to the true source positions and orientations. It turns out that in both MCMV and LCMV when spatial locations are set, optimal orientations that maximize the localizer can be found up to their sign as solutions of known eigenvalue problems. Thus, orientations can be excluded from the search for the maxima. For this reason, in the expressions below we assume source orientations to be known.

In a general case of oscillatory activity, an unbiased *MPZ* (that is, multi-source pseudo-Z) localizer is defined as follows:

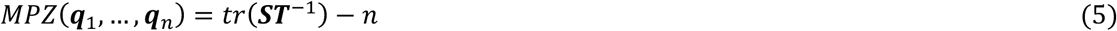

where (*n × n*) matrices ***S,T*** are defined as ***S*** = ***H***^*T*^***R***^−1^***H*** and ***T*** = ***H***^*T*^***NR***^−1^***H*** and ***N*** is the (*M × M*) noise covariance matrix. Note that noise should include all sources of electromagnetic field except the sources of interest targeted by the beamformer. Thus, the noise covariance includes instrumental and environmental noise as well as background brain activity. In the single source LCMV case *MPZ* is reduced to a well-known expression for pseudo-Z (denoted as 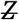) up to a constant offset equal to 1:

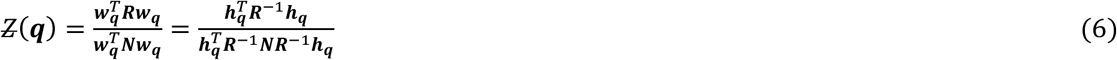

When dealing with so called evoked activity, that is time locked to a stimulus, significant increase in SNR and spatial resolution may be achieved by averaging over epochs, which eliminates oscillatory activity as well as noise that are not phase-locked to the stimulus. This leads to event-related (evoked) localizers utilizing epoch-averaged fields. The multi-source evoked response localizer (MER) is defined by expression

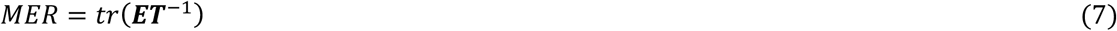

with (*n × n*) matrix ***E*** defined as 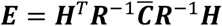, and 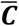 is the second moment of the averaged (“evoked”) field **〈*b*〉**:, 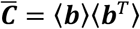. In a single source case (LCMV) this expression becomes

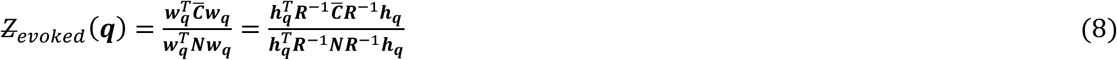

which is analogous to single-source evoked localizers introduced in (Robinson 2004; Cheyne et al. 2007).

In practice, maxima of the LCMV localizers are easily found by a brute force approach – that is, by calculating localizer values in each and every brain location. In the MCMV case this can’t be done (at least, with a reasonable computational cost) due to high dimensions of the parameter space, and accordingly, approximate iterative procedures are applied (Moiseev et al. 2011, Moiseev and Herdman 2013, Herdman et al. 2018). Those procedures start with a single-source LCMV beamformer as a first step. In subsequent steps, the beamformer order is incremented, and in, the simplest variant, already localized sources remain fixed and only the next source is searched by scanning the whole brain volume. In a more complicated and computationally intensive algorithm, the previously found sources are corrected in a secondary iteration loop (Herdman et al. 2018). In this work, the first (simpler) approach was used.

An important requirement for accurate localization is having a good estimate of the noise covariance ***N***. In task-based studies including the ones presented here, it can be reliably estimated during baseline, stimuli-free intervals, where no task-related activity is present. In resting state studies there is not such an option, and usually diagonal noise (like in this work) or some other substitute for the true noise covariance is used. Locations of interest are also typically defined *a priori* and do not need to be searched for. Still, without a true noise covariance source orientations are determined inaccurately, and should never be regarded as approximating the actual ones.

### Augmented pairwise MCMV (APW-MCMV)

In this study, we also introduce “augmented pairwise MCMV” (APW-MCMV) and test its performance relative to LCMV and traditional MCMV for resting state MEG connectivity analysis. In resting state network analyses connectivity is often estimated for all possible pairs of locations of interest (PW-MCMV). A straightforward way to avoid linear mixing, in this case, is to apply 2^nd^ order MCMV beamformer to the target pair. However, this only eliminates direct leakage from one source to another. If either of sources in the pair leaks into a location having strong connection with the other source, *indirect* leakage to a second source will occur, leading to spurious connectivity. Most likely such indirect leakage should happen via areas adjacent to the sources in the pair. This problem can be avoided by including all potential “conductors” of indirect leakage into the beamformer, but this approach has its limitations because besides computational considerations increasing beamformer order adversely affects the SNR of the filter.

Accordingly, we introduce a compromise solution called APW-MCMV, which proceeds as follows. First, connectivity for each pair of sources is estimated using standard pairwise MCMV, and statistically insignificant connections are discarded as described later. Then for every significant pair, we look for neighboring sources within a radius of 4 cm of each of the sources in the pair which do have significant connections. If those exist, up to two neighbors of each source in the pair with a maximal total number of connections are included in the beamformer. When estimating activity from the source pairs, the contribution from their neighbors are taken out. Thus, the maximum MCMV order for APW-MCMV calculation is 6. This higher order beamformer reconstruction is repeated for every significant pair, and new connectivity estimates are made which are now much less likely to be affected by indirect leakage.

### Connectivity measures

For task-related studies, we used coherence and phase locking values to compare the performance of LCMV and MCMV, as metrics sensitive to phase information are most popular in task-based electrophysiological connectivity analyses. To this end, reconstructed brain signals were estimated using beamformer weights derived from broad-band sensor time series. Time-frequency analysis was performed using a multi-taper strategy for decomposing the signal and estimating connectivity in a spectrally-resolved manner (Oostenveld, Fries, Maris, & Schoffelen, 2011). The multi-taper’s frequency-wide smoothing was set to 2 Hz. In the connectivity analyses, we used coherence and phase locking value (PLV) as they capture instantaneous synchrony. Their expressions can be found in Appendix A.

For resting state studies, we compared connectivity from LCMV, PW-MCMV and APW-MCMV using downsampled envelope correlations, due to their popularity in resting state MEG analyses and due to their previously demonstrated reliability (Colclough et al., 2016). At rest, alpha band oscillators are known to be located in occipital and parietal areas (Goldman, Stern, Engel, Cohen, & Cohen, 2002), thus resting analyses were carried out at the alpha band as this provides a physiological reference when interpreting connectivity results. Sensor data were band-pass filtered to the 8-12 Hz frequency range. To explore the possibility of using MCMV as an alternative to more conservative methods based on discarding 0-lag connections, we compared the results with the symmetric orthogonalization (SO) technique applied to LCMV reconstructed time courses (Colclough et al., 2015). For the same data, we reconstructed the time courses using LCMV, PW-MCMV and APW-MCMV, and LCMV with SO. In all four cases, amplitude envelope correlations were used to estimate connectivity. The envelopes of the signals were computed by taking the absolute values of the analytic Hilbert transform of the signals and then low-pass filtering to 0.5 Hz.

### Surrogate data

To create sensor- and source-level surrogate noise data, two types of surrogate data were used. First, we generated sensor-level surrogate datasets based on subjects resting state data, preserving the data covariance matrix but destroying all true connectivity within this data. This was done by randomizing phases of the principal components of the sensor data covariance matrix in the frequency domain. Datasets created this way were used in the task-related simulations for construction of the brain noise background, and in the resting state for the surrogate analyses.

Second, we generated source-level Gaussian surrogate datasets based on source reconstructed time courses of the real resting state data. The goal was to have surrogate source signals with time structure similar to real source signals but having no pairwise connectivity among them. This was done by fitting an autoregressive (AR) model of order one to the reconstructed time courses of real data, then using this model to generate random Gaussian source time courses with the same temporal smoothness. This was used to determine statistically significant pairwise envelope correlations in resting state analyses.

### Significance testing

To find statistically significant envelope correlations we followed a method suggested in Colclough et al., (2015). Briefly, the correlations were first mapped to [-∞,∞] interval using Fisher transformation, then z-scores of the results with zero mean and unit variance were estimated. To calculate the z-scores, we found standard deviations of the transformed correlations of the surrogate connectivity-free data generated using the AR model. The z-scores were then transformed to p-values under Gaussianity assumption, and those were corrected for multiple comparisons using a False Discovery Rate (FDR) threshold equal to 0.05.

### Forward modeling

For all source reconstructions, forward solutions were constructed using a single shell head model describing the inner skull boundary (Nolte, 2003). In simulation analyses, the sensor level signal to noise ratio (SNR) was defined as the square root of the ratio of the total average power of all sensors of the pure signal dataset to the total average power of the noise dataset.

#### Task related analyses

##### Simulation I: sinusoidal signals

To assess the effects of signal leakage we generated four white Gaussian random signals and added to them sinusoidal activity at two distinct frequencies. Signals were generated as:

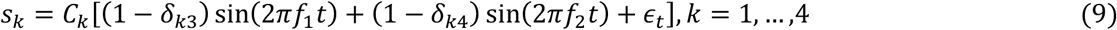

Here *f*_1_=10 Hz, *f*_2_=35 Hz, *t*is time in seconds, ε is a standard independent Gaussian zero mean random number chosen independently for each time point: ε_*t*_~*N*(0,1), and *δ*_*ij*_ is Kronecker delta. For each *k* constant *C*_*k*_ was chosen so as to obtain signal RMS equal to 10 nA. Thus, coherence between all sources at 10 Hz was close to 1 except for source 3. Similarly, coherence at 35 Hz for all sources was close to 1 except for source 4. The source signals were projected to the sensors using the lead fields from the ECoG simulation explained below. Surrogate data from a resting state MEG recording was added at the sensor level to model connectivity-free brain noise with an SNR of 2.4. Broad-band covariance (1 – 50 Hz) was used to calculate beamformer weights. The aim of this simple simulation was to demonstrate the effects of signal leakage and source cancellation in the LCMV case and compare the results with the MCMV case.

##### Simulation II: ECoG signals

Electrocorticogram (ECoG) data were used to simulate realistic brain signals, with the same complexity as signals measured with MEG. Data were obtained from an open ECoG database (Miller, 2016), and as requested by the author we reproduce the ethics statement:

> The ECoG patient participated in a purely voluntary manner, after providing informed written consent, under experimental protocols approved by the Institutional Review Board of the University of Washington (#12193). The patient data was anonymized according to IRB protocol, in accordance with HIPAA mandate. These data originally appeared in the manuscript *“Human Motor Cortical Activity Is Selectively Phase-Entrained on Underlying Rhythms”* published in (Miller et al., 2012).

ECoG time series collected from a pre-surgical patient flexing his thumb were used as ground truth. Thumb position was recorded using a 5 degrees of freedom data-glove sensor, thus trials were time locked with the movement onset. Four electrodes with the highest connectivity were selected. The lead fields for voxels closest to the actual electrode position were calculated and data was back projected to the sensors (Fig. 1A). Surrogate sensor data was added at the sensor level to model brain noise, and beamformer weights were calculated using a broad-band covariance (1 – 50 Hz). Different levels of noise were added to measure the beamformers performance as a function of SNR.

**Figure 1.**
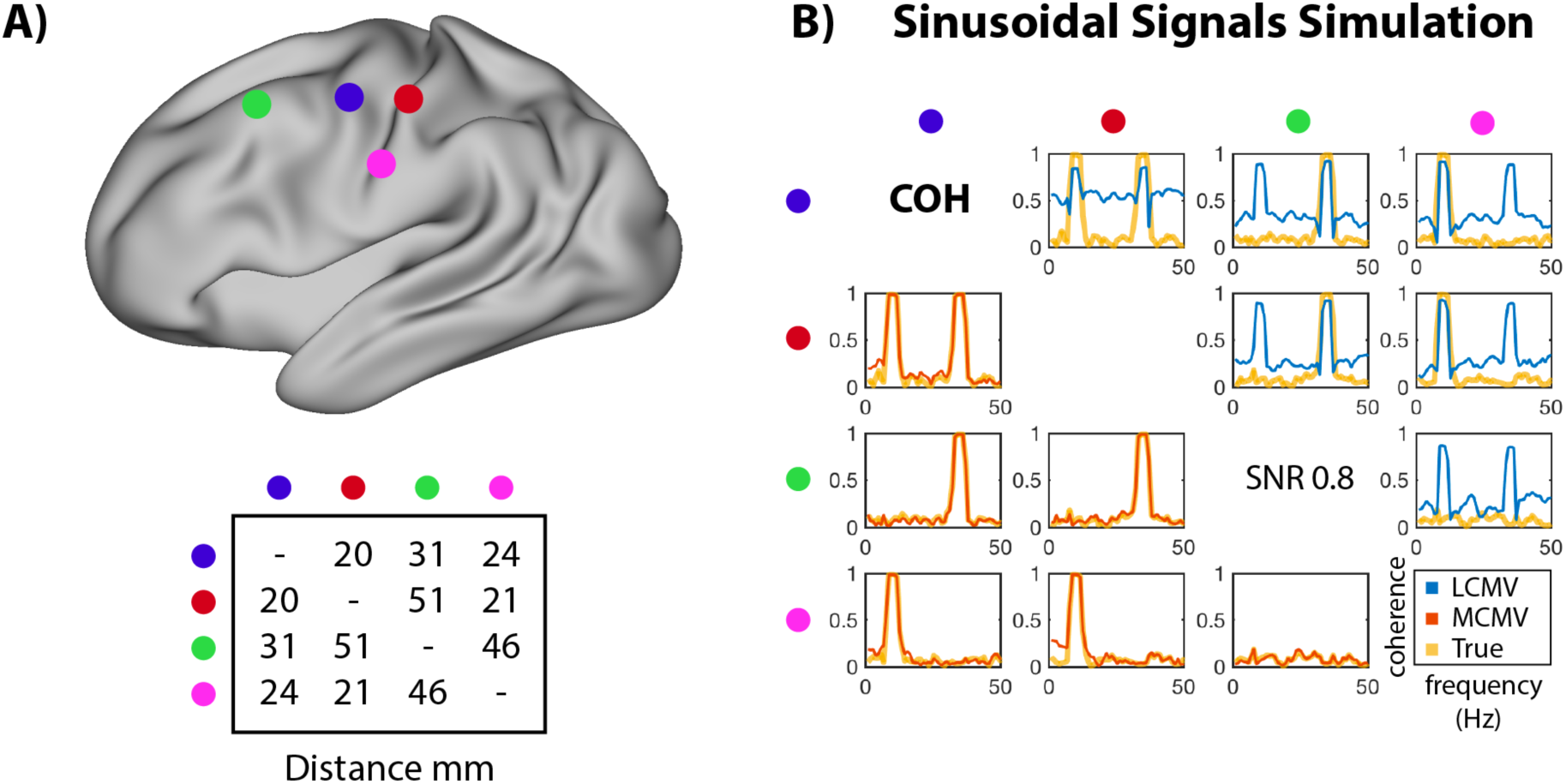
A) Projected locations of four sources and inter-source distances for sources used in Sinusoidal and ECoG simulations. B) Coherence between signals reconstructed using LCMV (upper triangle -in blue) and MCMV (lower triangle -in red); the true coherence is shown in yellow.

##### Real data: steady-state foveal entrainment

MCMV is very well suited for bilateral steady-state brain entrainment in which bilateral sources oscillate at the same phase and frequency producing strong correlations. While MCMV has no problems with source correlations, LCMV performance is likely to degrade due to the source cancellation effect, and signal leakage should overestimate connectivity in frequencies outside the entrainment.

Data were recorded from four subjects using a 275-channel whole-head MEG system (CTF Systems Inc., Coquitlam, Canada) at a sampling rate of 1200Hz. The study was approved by the ethics board of Simon Fraser University and all participants consented for the study. The stimulus consisted of a checkerboard circle of 4 degrees alternating black and white at 12 Hz for a period of 3 seconds, followed by an intertrial interval of 3 seconds with a fixation cross.

The first stimulus response (FSR) occurs at 6 Hz, half of the entrainment rhythm, from 0.1 to 0.3 sec, and this strong response was used to generate the second moments averagematrix 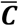 for the MER localizer. The signal covariance matrix ***R*** was computed from 0.1 to 3 seconds period, and the noise covariance ***N*** from 2.8 to 0 seconds period. A 5 mm grid was used to find two MER maxima. Then, coherence between the two peaks was estimated between 4 and 45 Hz.

#### Resting-state analyses

The data used in the following resting state analyses were obtained from the Human Connectome Project (HCP) and the data were preprocessed as described previously (Larson-Prior et al., 2013). All analyses were performed with 76 sources representing each cortical region in the Automatic Anatomical Labelling (AAL, Tzourio-Mazoyer, et al., 2002) atlas, and envelope correlation was computed between the source pairs in the alpha band (8 – 12 Hz). These analysis choices were selected as they reflect popular approaches in the current literature. Sources were reconstructed using an LCMV, a PW-MCMV – two sources reconstructed for each pair-, and an APW-MCMV –where up to four additional sources were included in the beamformer as explained above. In addition, SO was applied to LCMV-reconstructed sources to remove 0-lag interactions. In total, connectivity was estimated with four methods: LCMV, PW-MCMV, APW-MCMV and SO

##### Single subject connectivity analysis

Using resting state data collected from a single subject as an example, we compared the four different approaches for estimating functional connectivity using envelope correlations in the real data. Additionally, we performed the same analyses for the simulated data where the ground truth was known. The latter was generated by projecting SO-reconstructed time series back to the sensors and adding surrogate connectivity-free resting state noise background to obtain sensor-level SNR = 2.

##### Group-level connectivity analysis

Connectivity was computed across all the 89 subjects present in the HCP dataset. A fixed effect approach was used to find significant connections and multiple-comparisons corrected using FDR. Then, to assess if the effects found might be driven by signal leakage, the same connectivity analysis was performed using the connectivity-free surrogate data generated using the actual resting state data for the respective subject. Any significant results found in the surrogate analysis would likely indicate spurious connectivity, because even if at the individual level some connections might be significant by chance, at the group level only consistently strong leakage-driven connections would survive.

## Results

### (a) Task related analyses

#### Simulation I: sinusoidal signals

We first contrasted the performance of LCMV and MCMV in task-related coherence analysis using simple simulations of sinusoidal signals (Fig. 1B). For LCMV, the coherence at 10 and 35 Hz is very high for all pairs even though some of the true signals were not coupled at these frequencies. Moreover, LCMV coherence is high across all the frequency spectrum while the original signals were only coupled at two discrete frequencies. This signal-leakage spurious connectivity is not present in the MCMV case which is capable of estimating the original connectivity. In addition, due to signal cancellation occurring in the LCMV case, the maximum coherence reached at 10 and 35 Hz is reduced compared to MCMV and to its true value.

#### Simulation II: ECoG signals

We then contrasted the performance of LCMV and MCMV for coherence and PLV analyses using more realistic simulated MEG data by using ECoG data (Fig. 2A), and varied the sensor-level SNR by scaling the added surrogate background MEG data. At an SNR of 2.5, connectivity was estimated using coherence and PLV (Fig. 2B). The most striking difference between LCMV and MCMV is seen with the motor and sensory source pair (blue and red in Fig. 1A), where LCMV estimates a very high spurious connectivity across all frequencies except for those below 5 Hz, where the estimated connectivity is smaller than the true one. This is caused by source cancellation which is getting stronger with the SNR, and the frequency-specific SNR at low frequencies is the highest. At SNR = 2.5, MCMV is capable of capturing the true connectivity with both coherence and PLV, although PLV results are noisier. The latter effect might be explained by the nature of the PLV measure which does not take instantaneous signal amplitude into account thus artificially elevating contributions of small amplitude noise components to the result.

**Figure 2.**
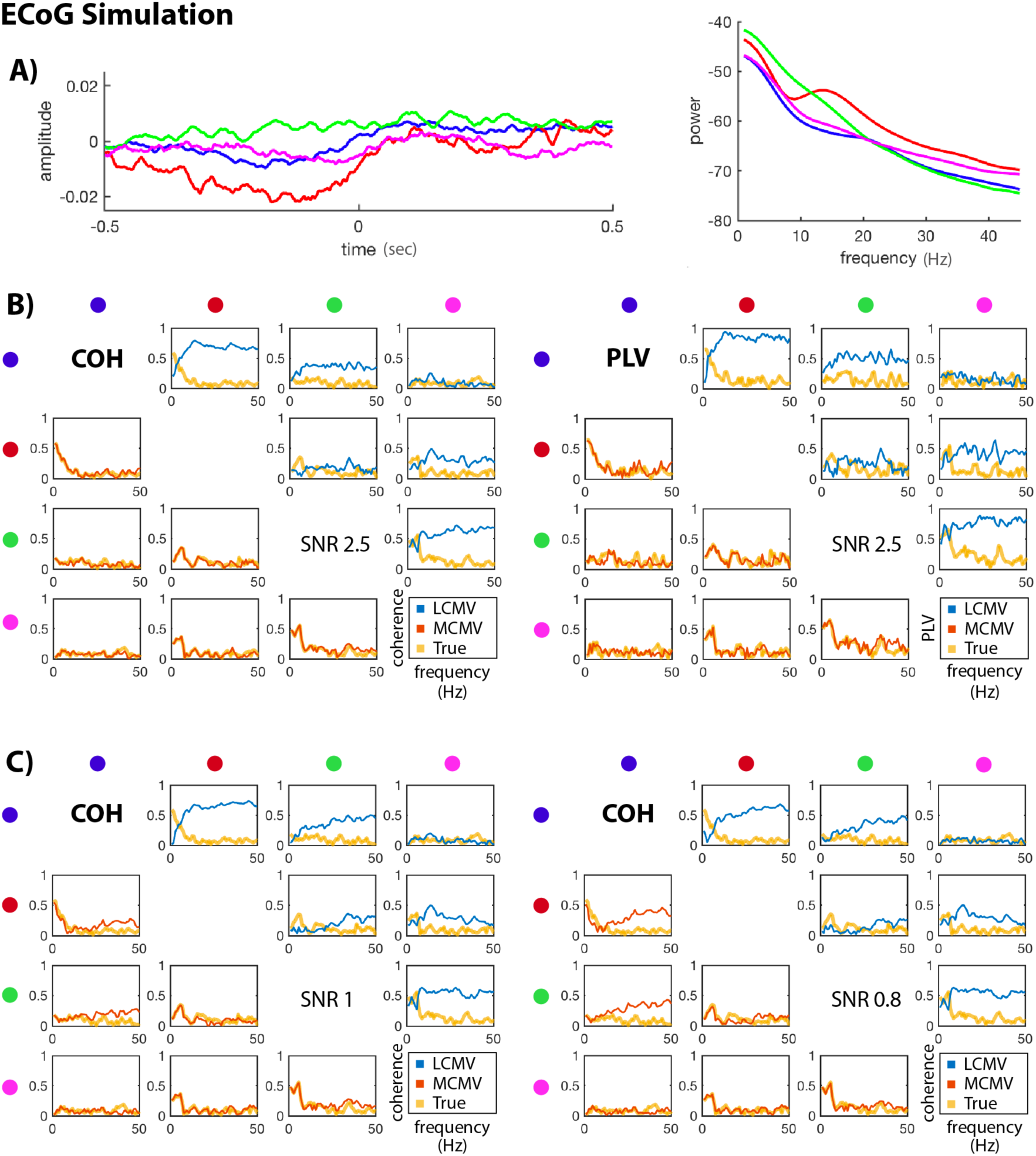
A) ECoG epoch-averaged time series used in the simulations, and their power spectrum. B) Coherence and PLV of the reconstructed time courses for SNR = 2.5 C) Coherence of reconstructed time courses for SNR = 1 (left) and SNR = 0.8 (right).

At SNRs below 1, the performance generally degrades for both beamformers, this effect being more pronounced at higher frequencies. This happens due to two reasons. First, the beamformer weights constructed with broadband covariance are better optimized for low frequencies which carry most of the power. Second, at the source level, frequency-specific SNR of ECoG signals is lower. In the MCMV case, the most degradation of the performance is observed for pairs with smaller inter-source distances (such as the blue labeled source with the red or green in Fig.2C). Due to the finite resolution of the spatial filter, the SNR of the nth-source is affected by the null placed on another source if it is close by.

#### Real data: Steady-state Foveal entrainment

Next, we contrasted the performance of LCMV and MCMV using real MEG data collected during a foveal entrainment paradigm, which was chosen because the relevant neuroanatomy and neurophysiology are relatively well characterized. Brain entrainment occurs when a stimulus is repetitively presented at a certain frequency. In this study, visual areas representing the fovea were entrained at 12 Hz by presenting a circular grating at about 4-5 degrees in the retina and contrast alternating 6 times per seconds.

In MCMV source localization employed here (Moiseev et al., 2011), in the first iteration maxima of the MER localizer Eq. (8) coincide with those of LCMV’s evoked pseudo-Z Eq. (9). Importantly, due to source cancellation effect, there is only a single peak corresponding to the strongest source. The other source becomes visible only at the 2^nd^ iteration, when the activity of the 1^st^ source is “nulled” (see brain plots in Fig. 3) and correlated interference is removed.

**Figure 3.**
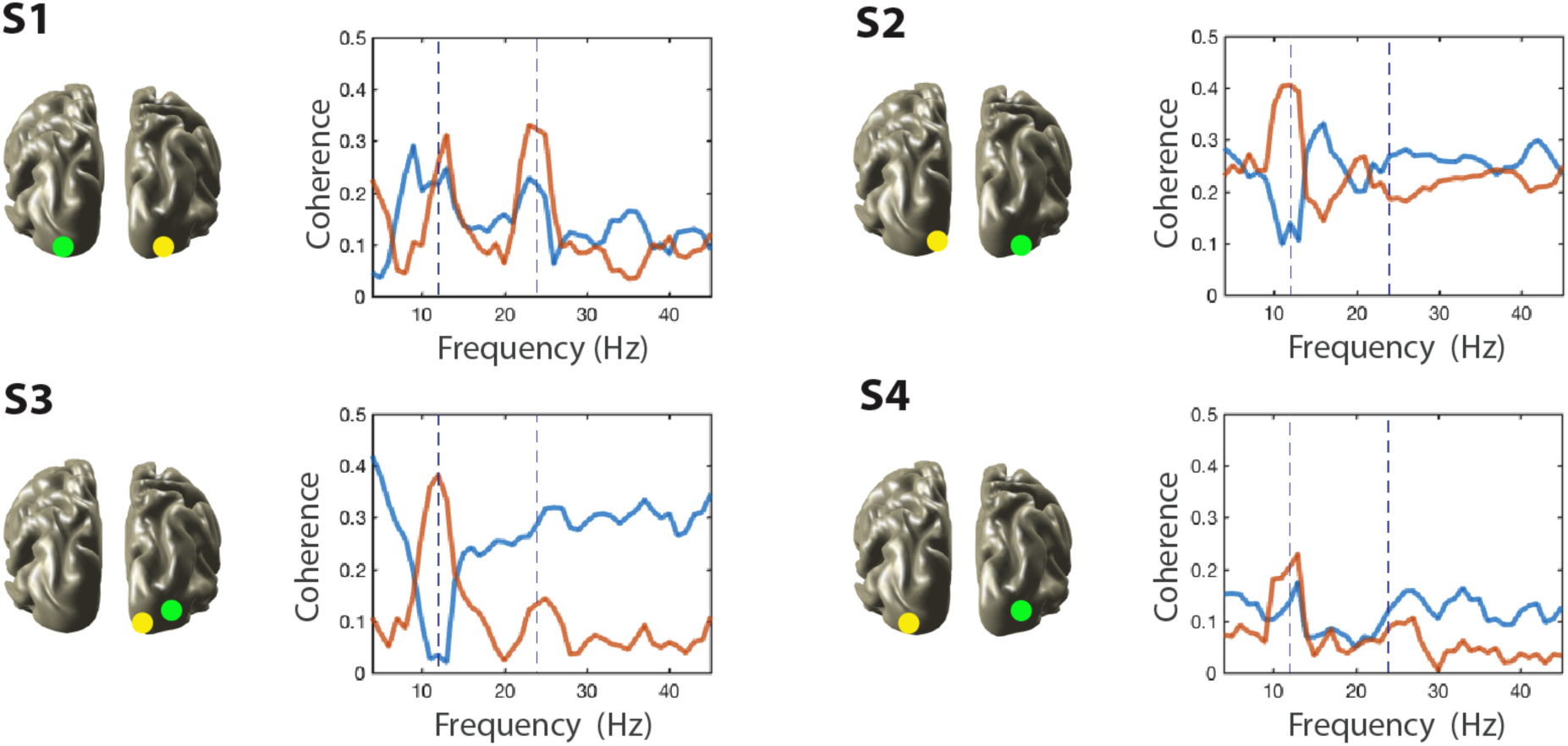
Visual entrainment of a visual stimuli oscillating at 12 Hz. Four subjects S1 – S4 are presented. Projections of locations of peaks 1 and 2 on the cortex surface are shown by yellow and green circles, respectively. Coherence between the two peaks found was estimated in [4, 45] Hz band using LCMV (blue) and MCMV (orange) reconstructed time courses.

The first two found MER peaks corresponded to either the foveal representation or the V4 visual area sensitive to motion. Coherence between those was computed using broadband (1 – 50 Hz) beamformer weights for source reconstruction (Fig. 3 connectivity plots); for LCMV estimates, the 2^nd^ source location found by MCMV was used. As can be clearly seen in Fig. 3, LCMV results generally do not show pronounced isolated peaks at the entrainment frequency or its harmonics. Instead, they often reflect strong source cancelation at 12 Hz (subjects S2, S3). In contrast, MCMV produced a distinct peak at 12 Hz in all cases and also peaks at 24 Hz for subjects S1 and S3. Also, LCMV generally showed increased broad band coherence compared to MCMV which is explained by uncontrolled leakage.

### Resting-state analyses

#### (i) Single subject connectivity analysis

To compare the performance of LCMV, PW-MCMV and APW-MCMV for resting state MEG connectivity we first contrasted the results in a single subject analysis. First, we estimated connectivity in the actual real data using four methods – namely LCMV, PW-MCMV, APW-MCMV and symmetrically orthogonalized LCMV time courses (SO). Then, we used the time courses obtained by SO to construct a simulated dataset where the ground truth connectivity was known. Specifically, those time courses were projected back to the sensors and connections-free surrogate brain noise data was added at the sensor level. Thus, significant connections discovered in this data which did not exist in the original SO time series must be spurious, and would suggest that the same significant connections in the real data were also spurious.

Results for single subject analysis of resting MEG connectivity are presented in Fig. 4A. Alpha band connectivity at rest was expected to be driven in large part by connections involving occipital and parietal areas (Goldman et al., 2002). Clearly, the SO results confirm these expectations. On the contrary, LCMV-based estimates demonstrate strong connections in inferior-frontotemporal and midline areas in addition to parieto-occipital areas. PW-MCMV results are much closer to SO results but still show a few significant connections in inferior-frontotemporal areas. APW-MCMV further corrects the pair-wise approach by discarding most of the connections located anteriorly. With real data it is not possible to be certain if connections are real, or for that matter, if SO results are free from spurious connections.

**Figure 4.**
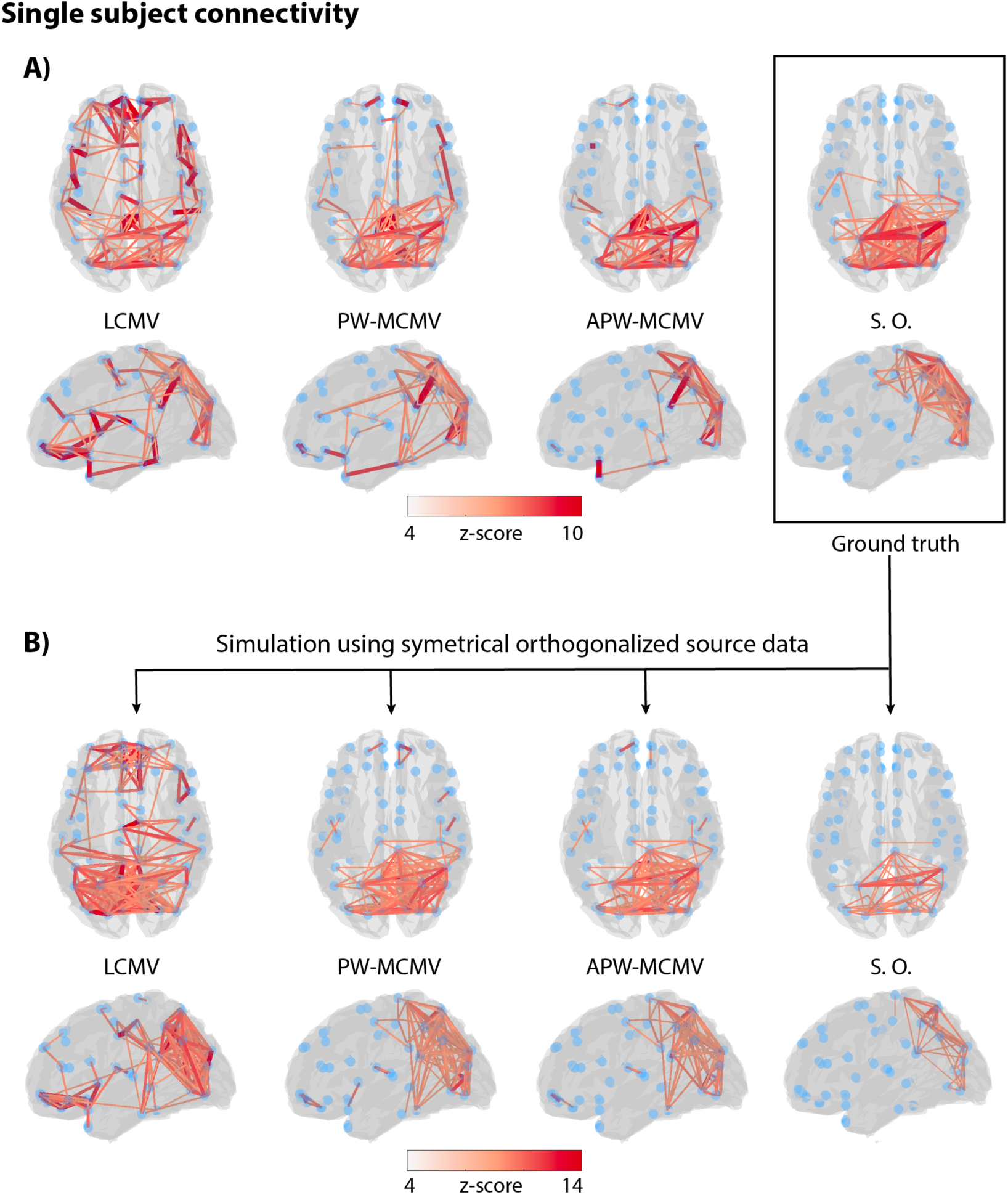
Single subject resting state connectivity analysis. A) Significant alpha-band amplitude envelope correlations in the real data obtained with LCMV, PW-MCMV, APW-MCMV and SO. B) Same analyses applied to the simulated resting state data with the “ground truth” connectivity corresponding to the right-most panel of (A)

To identify spurious connections, we simulated data using the SO reconstructed time courses. Thus, the “true” functional connectivity graph was known, and corresponded to the right-most plot in Fig. 4A. The simulation results are depicted in Fig. 4B. Pure LCMV-based connectivity estimates still shows abundant anterior, fronto-temporal and other connections which we can now confidently characterize as spurious. Conversely, the SO approach which seemed the most reliable in the real data turned out to be conservative with numerous true connections were discarded, even though the “true” source time courses were already orthogonalized and therefore ‘true’ connectivity should not be discarded. PW-MCMV and especially APW-MCMV results were the ones most close to the ground truth, with APW-MCMV having less spurious connectivity.

#### (ii) Group-level connectivity analysis

To further contrast the performance of LCMV, PW-MCMV, APW-MCMV and SO for resting MEG connectivity, analyses described in the previous section were applied to each of the 89 subjects in the HCP dataset, then results were further corrected for multiple comparisons at 0.05 FDR level (Fig. 5). In general, group-level results follow the pattern observed in the single subject case, with LCMV demonstrating lots of potentially spurious connections and SO being the most conservative.

**Figure 5.**
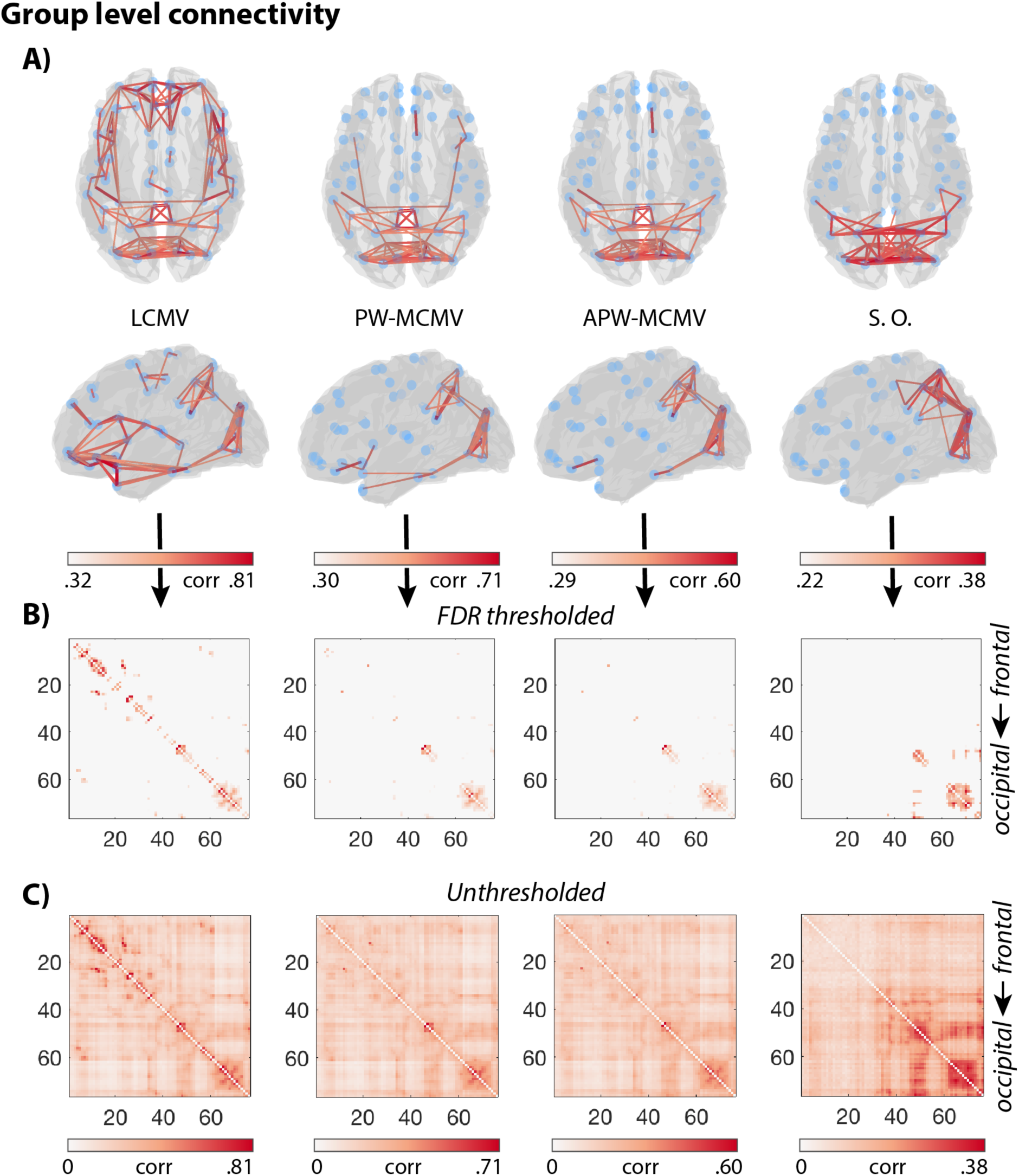
Group-level envelope correlation analysis. A) Significant connections FDR corrected plotted in brain space. B) Connectivity matrices thresholded with only significant connections. C) Raw connectivity matrices depicting the spatial connectivity pattern. For visualization purposes, the correlation values were used instead of the z-scores. Connections closer to the diagonal represent spatially closer source pairs.

#### (iii) Surrogate data group-level analysis

To contrast the susceptibility of each of the compared methods to spurious connectivity, we performed a connectivity analysis of ‘null’ surrogated data. Compared to the single subject analysis, however, these seemingly spurious connections in the group level could be real as the ground truth is unknown. To reveal connections that are guaranteed to be false we performed the same group level connectivity analysis with each subject’s data replaced with corresponding connectivity-free surrogate sensor data. After that, no significant connections should have been found on a group level. This was indeed the case with SO, while with LCMV a great number of connections were found significant in inferior frontotemporal and midline areas as in the previous single subject analysis. In the MCMV cases the number of “significant” spurious connections were very small, particularly with APW-MCMV with only 3 false positives (Fig. 6A). Still, it can be noted that while SO spatial connectivity lacked any discernable pattern (Fig. 6B), for MCMVs connectivity matrices there is a clear spatial pattern. This pattern is very pronounced in LCMV, giving rise to many false positives. The pattern increasingly improves and fades out with PW-and APW-MCMV, albeit do not completely disappear.

**Figure 6.**
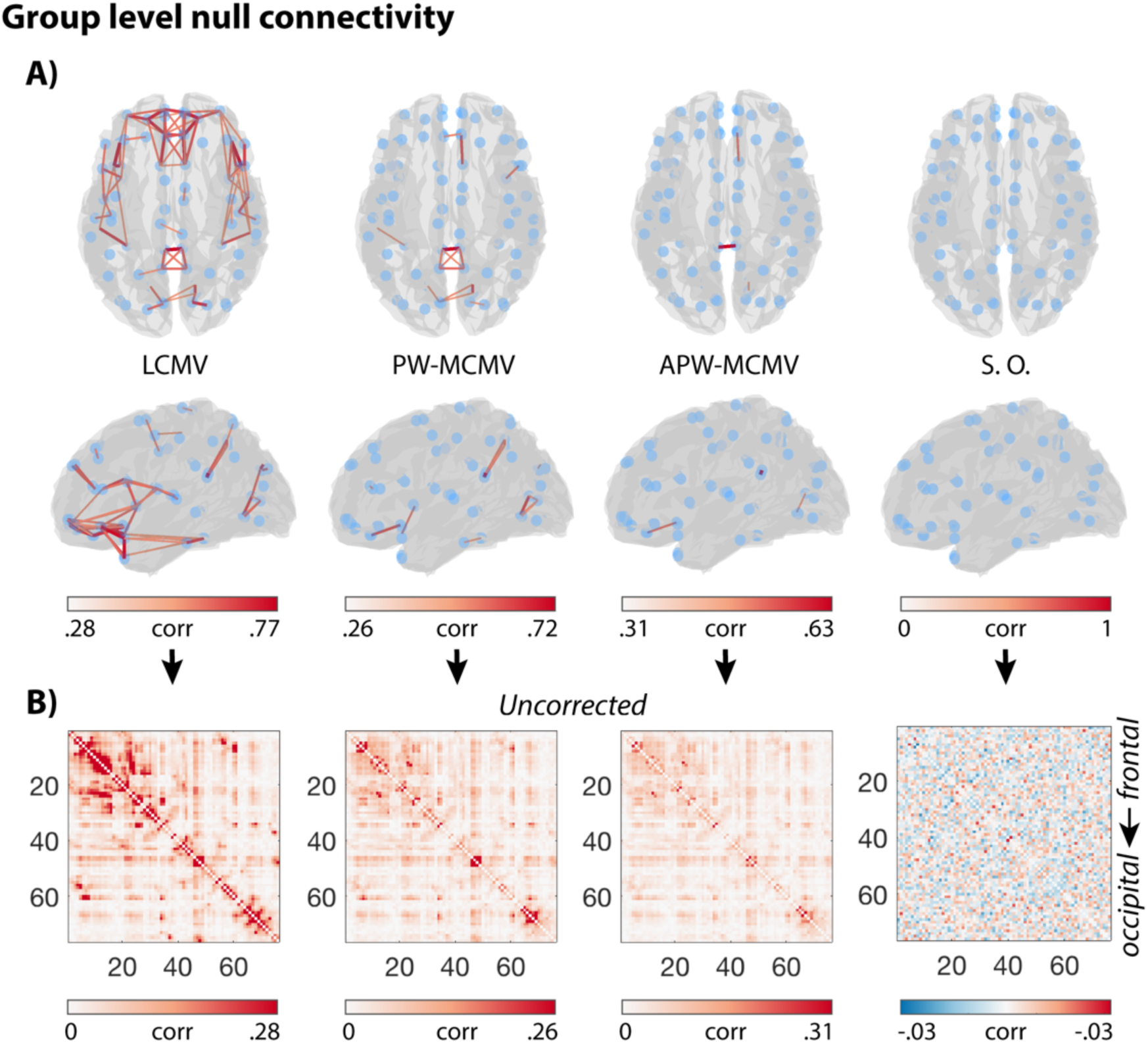
Connectivity for each subject estimated using surrogate sensor data with the phase randomized. A) Significant connections on a group level for surrogate data. B) Correlation matrices with the upper color threshold being the smallest significant correlation value. Connections closer the diagonal represent spatially closer source pairs.

## Discussion

In this study, we demonstrate the advantages of using MCMV method when estimating functional connectivity in the brain based on beamformer source reconstruction. We show using real and simulated data that MCMV outperforms LCMV by better suppressing signal leakage and coherence signal cancellation in task conditions using inter-regional phase synchronization and coherence analysis, and in resting state for mapping inter-regional amplitude correlations. In addition, for resting state analyses we introduce an APW-MCMV approach which further improves signal leakage correction compared to PW-MCMV.

To a large degree, MCMV prevents signal leakage, which is the main cause of spurious connectivity found by a traditional LCMV beamformer. An alternative method to achieve this goal is to discard 0-lag connections. This can be done in many different ways but in this work we chose symmetrical orthogonalization approach described in Colclough et al. (2015) for comparisons, since the latter shows a good performance and provides a strong control for false connections. Especially for the resting state scenarios, we wanted to investigate if MCMV could deliver similar results but without limitations imposed by that method. In particular, we wanted to be able to address situations when connections with short propagation times are important and should not be discarded, or when one needs to look at finer time scales than amplitude time courses would allow, so that other connectivity measures, such as PLV or coherence, should be applied.

We focused on connectivity analyses of task-related activity and resting state activity, which are the two most common scenarios encountered in basic and clinical research. There are differences in how beamformer reconstruction is performed in each case. In task-related paradigms typically there are few sources of activity, but locations of those are not known in advance. Consequently, beamformer reconstruction involves a source localization step. Importantly in this case, noise covariance can be properly estimated using control intervals, and the signal to noise ratio (SNR) may be high enough due to trial averaging. By applying MCMV localizers (Moiseev et al., 2011), it becomes possible to accurately find source positions and orientations even for highly correlated sources, as demonstrated in our foveal entrainment experiment. In the resting state analyses source locations are usually selected *a priori*, however the number of sources is large and SNR is close to zero. Moreover, there is no way to estimate the noise covariance, because it is impossible to separate an elementary resting state brain source “signal” from the rest of the background brain activity. Consequently, it is impossible to calculate source orientations accurately. Both factors result in poorer source reconstruction and less reliable connectivity estimates.

We demonstrated here that in both task and resting state scenarios MCMV outperformed LCMV by greatly reducing, albeit not completely discarding, spurious connectivity, despite the fact that 0-lag connections were included. For the resting-state functional connectivity, our *augmented pairwise-MCMV* approach was shown to further reduce spurious connections compared to simple *pairwise*-MCMV.

The ECoG simulation data were used to provide some insight into how LCMV and MCMV performance are affected by SNR. It is known that increasing beamformer order leads to a gradual decrease in SNR of the reconstructed time series, although theoretically this should become noticeable only if the order is comparable to the total number of sensors in the array. Our simulation results showed that sometimes such effects can be felt much earlier. Specifically, due to finite spatial resolution, the nulls imposed by an MCMV beamformer are not points but rather areas of certain size in the signal space. If the sources are sufficiently close to each other, the nulls imposed on neighboring sources overlap with the target source. As a result, MCMV weights become less robust and very “fine-tuned” to the interference. In our particular case, such overlap did not cause significant problems when the SNR was high. In low SNR cases when a broadband covariance was used, beamformer weights still worked well for lower frequencies where most of the power was concentrated. However, these weights turned out to be not that optimal for higher frequencies, resulting in MCMV reporting spurious connectivity there (see Fig. 2C). This effect was observed for pairs involving closer sources (blue and red, blue and green in Fig. 2C). When frequency-specific narrow band covariance was used for weights construction, this spurious connectivity disappeared. Accordingly, using a narrow-band covariance (that is, frequency-dependent weights) whenever possible seems advantageous because in practice neither signals nor noise have flat power spectra. It is also interesting to point out how LCMV filter inverts the true signal coherence (Fig. 2B, 2C) yielding completely distorted frequency dependence. In this case, source cancellation played a major role. The cancellation was most pronounced at the low end of the spectrum where both the true coupling and the SNR were high. As a result, reported coherence was minimal. At higher frequencies, the source signals canceled less, and estimated coherence grew due to strong uncontrolled leakage.

The human steady-state foveal entrainment MEG study provided a good example of a situation where LCMV approach fails due to both signal leakage and source cancellation occurring for strongly correlated activity. At the same time, methods of leakage correction that discard the instantaneous interaction between two signals would be obviously unsuitable here. In this experiment, MCMV revealed connectivity in areas associated with foveal representation, whereas LCMV connectivity was smeared across the visual cortices. Based on this, we believe that future studies could benefit from using MCMV to estimate highly synchronous activity occurring when separate brain areas are simultaneously paced by the same stimulus.

In our resting-state analyses, we compared connectivity estimates obtained with LCMV, MCMVs and SO approaches at a single-subject and group levels, for real and surrogate data. We used the SO approach as a conservative reference method with a strong control for spurious connectivity due to signal mixing at the expense of discarding zero-lag connections.

In the single-subject case, we found that LCMV greatly overestimates connectivity in inferior frontotemporal and midline areas. In contrast, with orthogonalized signals the only significant connections were located in posterior-parietal and occipital areas, where alpha generators are expected (Goldman et al., 2002). Pairwise (two-source) MCMV mostly eliminated spurious couplings reported by LCMV, however, it still reported more connections than SO. The APW-MCMV results were close to the SO ones. At the same time, the simulation where the ground truth connectivity was known, showed that even at relatively high SNR = 2, the SO method was conservative discarding true connections but keeping only true ones, while MCMV-based approaches preserved those, at the expense of showing more false positives. This illustrates a well-known tradeoff between type I and type II statistical errors where in our case the SO method showed a strong control over type I error while allowing large type II error (i.e. rejecting the true positives). It is worth noting though that in practical situations controlling for type II error is as important or even may be a priority. The MCMV-based approaches seem to suggest more balance between the two errors, and have an additional advantage of taking 0-lag connections into account.

The group level analyses confirmed the results found in the single-subject case. When the group analysis was performed with the subjects’ connectivity-free surrogate data, we found that the connectivity matrix for SO did not display any spatial pattern, as should be expected. LCMV and MCMVs had a clear spatial pattern, however, while in LCMV this pattern was strong enough to produce lots of false connections appearing as being significant, with PW-MCMV only a few such connections remained, and with APW-MCMV those were almost eliminated.

### Conclusion

In this comprehensive study, we investigated the advantages of MCMV spatial filters for analyses of noninvasive electrophysiological functional connectivity in the human brain. Using simulated and real data in task-related and resting state scenarios, we demonstrated how MCMV-based approaches address the problem of spurious connectivity arising from signal mixing and coherent source cancellation, without discarding connections with very short or zero-time delays. We also showed that MCMV, while strongly reducing spurious connections obtained with traditional LCMV reconstruction, provides better control over type II statistical error than more conservative methods based on completely discarding 0-lag interactions between the signals.

## APPENDIX A

PLV is a measure of phase synchrony between two signals calculated by averaging the phase difference 𝜃 between complex-valued narrow-band time series (Lachaux et al., 1999). For a pair of such signals *x* and *y* PLV is estimated according to the expression:

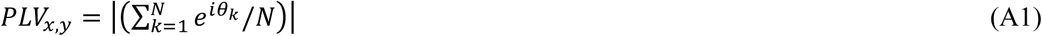

Here *N* is the number of epochs, and *θ_k_* is the phase difference between the signals in *k*-th epoch.

The coherence between two signals *x* and *y* is defined via their power spectra and cross spectra at specific frequency *f* by the formula:

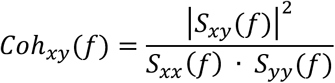

where *S*_*x,y*_ denotes the cross spectrum of the signals at frequency *f*.

## Acknowledgements

This study was supported by the Canadian Institutes of Health Research (http://www.cihr-irsc.gc.ca) CIHR grant to S. M. D. (MOP-136935) and NSERC grant to S. M. D. (RGPIN-435659).

